# Predicting environmental and ecological drivers of human population structure

**DOI:** 10.1101/2022.06.08.495166

**Authors:** Evlyn Pless, Anders M. Eckburg, Brenna M. Henn

## Abstract

Landscape, climate, and culture can all structure human populations, but few methods are designed to disentangle the importance of these many variables. We developed a machine learning method for identifying the variables which best explain migration rates, as measured by the coalescent-based program MAPS that uses shared identical by descent tracts to infer and extrapolate spatial migration across a region of interest. We applied our method to 30 human populations in eastern Africa with high density SNP array data. The remarkable diversity of ethnicities, languages, and environments in this region offers a unique opportunity to explore the variables that shape migration and genetic structure in humans. We explored more than twenty spatial variables relating to landscape, climate, and presence of tsetse flies (an important regional disease vector). The full model explained ~40% of variance in migration rate over the past 56 generations. Precipitation, minimum temperature of the coldest month, and altitude were the most important variables. Among the three groups of tsetse flies, the most important was the *fusca* group which is a vector for livestock trypanosomiasis. We also performed a selection scan on a subgroup of the populations who live in Ethiopia at relatively high altitudes. We did not identify well-known high-altitude genes, but we did find signatures of positive selection related to metabolism and disease. We conclude that environment has notably shaped the migration and adaptation of human populations in eastern Africa; the remaining variance in structure is likely due to cultural factors not captured in our model.

## Introduction

There is great interest in the mechanisms that determine gene flow and cause population structure, but there are computational challenges to disentangling the many factors at play. In humans, there is a variety of complex and often correlated environmental variables – land cover, climate, culture, and language – that all affect migration patterns and resulting population structure. In general, we know populations farther apart tend to exchange fewer migrants (“isolation by distance” (Wright 1943)); however, gene flow does not always scale proportionally with geographic distance. A mountainous region might limit gene flow between two human populations more than equally spaced populations at the same elevation, and the same idea applies for cultural barriers like language. There are a variety of landscape genetics tools designed to associate environmental and spatial variables with gene flow (Manel et al. 2003). In landscape genetics, it is often useful to depict migration routes and barriers through “resistance surfaces,” in which each pixel has a value representing the difficulty of migrating across this pixel. Least cost path or circuit theory (McRae et al. 2008) can then be used to calculate “resistance distances” between pairs of population or individuals through these surfaces; these effective distances can then be associated with genetic distances to find correlations. Another method is to model genetic connectivity directly from the environmental data, for example using a maximum likelihood approach (Bouyer, et al. 2015). Other options include inferring migration surfaces (the inverse of resistance surfaces) from genetic distance data (EEMS) (Petkova et al. 2016) or from identical by descent tracts shared between individuals (MAPS) (Al-Asadi et al. 2019); however, these models do not explicitly include environmental or other spatial variables.

Previously, we developed an approach to infer landscape connectivity by integrating genetic and environmental data (Pless et al. 2021). This approach offered some important advantages, in particular the ability to deal with 1) correlated predictor variables (environmental) and 2) flexible, non-linear relationships between the predictor variables and the response variable (genetic distance) at different regions. A disadvantage of our model was that it used pairwise summary statistics like F_ST_ (Rousset 1997) and Cavalli-Sforza & Edward’s chord distance (Cavalli-Sforza and Edwards 1967) as proxies for genetic distance; these statistics make limiting assumptions and can be biased by demography. Additionally, it required the model to be based on pairwise distances among assumed populations rather than on individuals. Although these features were necessary give the use case (clustered sampling of *Aedes aegypti* mosquitoes genotyped with 12 microsatellites), they made the model more difficult to use and interpret. **In this manuscript we adapt and improve our previous method for high resolution human genomic data. We take advantage of identical by descent tracts to infer a migration surface and find what spatial variables predict migration using random forest, in a new approach we call SPRUCE (Spatial Prediction using Random forest to Uncover Connectivity among Environments)**.

We applied SPRUCE to eastern Africa, a compelling case study for understanding human population structure and migration. Specifically, we focus on southwestern Ethiopia, Kenya and northern Tanzania – which have the densest collection of publicly available genome-wide data. This region is rich with ecological diversity ranging from low desert to high, forested mountains. Additionally, the history of human migration into the region is complex (Prendergast et al. 2019), contributing to the high diversity of ethnicities, languages, religions, and subsistence strategies found there today (Atkinson et al. 2021; López et al. 2021). Cattle herding was introduced from the Near East into this region by ~4,000 year ago, and Bantu-speaking agriculturalists from western Africa later expanded into the Kenya/Tanzania border region by ~2,000 years ago (Hildebrand and Grillo 2012; Skoglund et al. 2017). Local hunter gatherers populations have exhibited varied responses to these migrations, including reducing their range, moving, adopting farming with varying levels of gene flow, or co-existing with the pastoralists and agriculturalists (Gopalan et al. 2022).

After applying SPRUCE to this complex region, we found precipitation, altitude, and temperature play important roles in shaping human migration in east Africa across three time intervals ranging from approximately 300-2,000 years ago. Given the importance of environmental variables in determining human migration rate in eastern Africa, we predicted that populations have adapted to microclimates in the region, which we tested by running genome-wide selection scans on five populations from southwestern Ethiopia. Although these populations live within 70km of one another, they practice a range of subsistence strategies and likely face different pressures in terms of pathogen exposure and high elevation, which can cause cardiovascular disease and pregnancy complications.

## New Approaches

### Extending SPRUCE for use with high resolution genomic data

We improve our previously developed method for inferring landscape connectivity (Pless et al. 2021) for dense genomic data. The primary difference between the previous method and the improved version of SPRUCE is how migration rates are calculated. In the original method, pairwise genetic distances (e.g. F_ST_) among all pairs of populations was used as proxy for migration rates. Genetic distance was then used as the response variable in a random forest regression, where the predictor variables represented environmental and other spatial variables of interest. In the first iteration, the predictor variables’ values were calculated by taking the mean of the pixel values found along straight lines connecting each pairs of points; in subsequent iterations “least-cost path” lines were used to represent a more realistic migration path through the landscape.

In contrast, for SPRUCE we calculate migration based on shared genomic patterns among individuals, rather than by using summary statistics like FST. After processing and phasing the genomic data (Fig. 1a), the first step of SPRUCE is to find and count genomic regions which are shared identical by descent (IBD) across each pair of individuals in the dataset; these are also known as “long pairwise shared coalescence” segments (Fig. 1b) (Al-Asadi et al. 2019). The number of shared IBD segments of certain lengths contains useful information for calculating relatedness and inferring migration; e.g. two individuals who are identical across long regions of the genome must share a recent common ancestor. The coalescent-based program MAPS takes a matrix of the number of shared IBD segments of a certain length for a set of individuals, along with the geographic coordinates for those individuals. It outputs inferred migration rate at each coordinate (“deme”) across the studied region (Fig. 1c). The response variable in the SPRUCE random forest regression is inferred migration rate at each deme; the predictor variables are the deme-level values of environmental and other spatial variables. These variables of interest are available from free online datasets, such as CHELSA for worldwide climate data (Karger et al. 2017), and values can be extracted with open source tools such as gdal (GDAL/OGR contributors 2021) (Fig. 1d).

**Figure 1:**
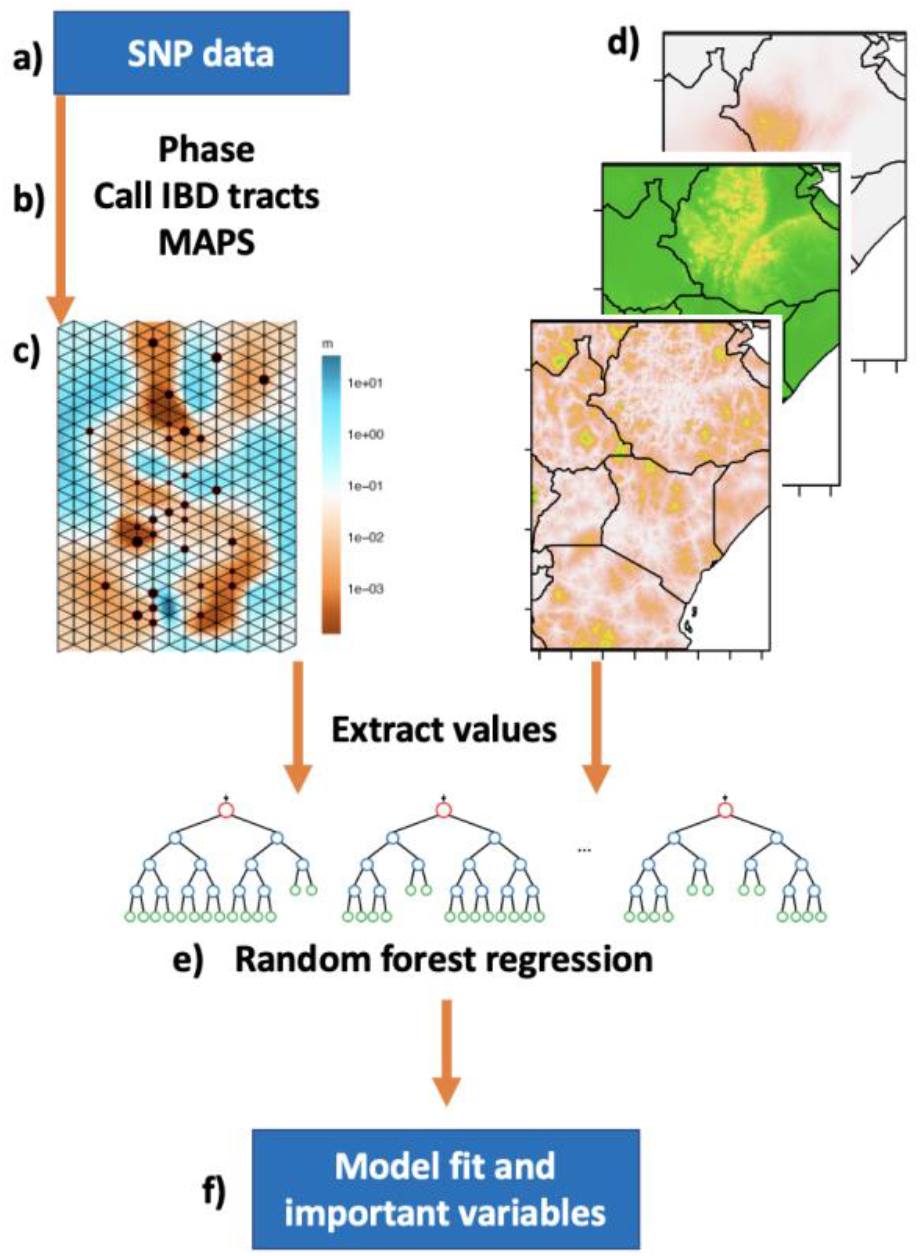
SPRUCE pipeline for integrating genetic and spatial data to infer ecological drivers of human migration. In short, the model uses a random forest framework to determine which spatial variables best predict migration, which is inferred from sharing of identical by descent tracts among individuals using the program MAPS (Al-Asadi et al. 2019).

Using a random forest framework (Fig. 1e), SPRUCE provides a straightforward approach for investigating which environmental variables contribute to migration rate, and how much variance these variables explain (Fig. 1f). In our test case, we showed that 25 variables explain ~40% percent of the variance in human migration in eastern Africa, and we validated the model by testing it on 30% of the data which was originally withheld. The SPRUCE approach introduced here is suitable for human datasets with genomic data that is dense enough to identify IBD regions (~400,000 SNPs or more). The model should be applied at a regional scale and within time frames where human migration can be inferred using IBD regions, and ideally the genetic samples are widely distributed throughout the region of interest. The model can accommodate a large number and wide range of predictor variables, including categorical data (e.g. language) and correlated datasets (e.g. elevation and temperature).

## Results

### SPRUCE Model

We developed a new methodology for determining the spatial variables that best predict human migration, as inferred from shared identical by descent tracts. Drawing genomic data from two sources (Gurdasani et al. 2015; Scheinfeldt et al. 2019), we generated a dataset of 517,383 SNPs for 492 individuals from 35 geographic locations within our region of interest (Fig. 2). After phasing the data and finding shared identical by descent tracts, we obtained spatial estimates of migration for three approximate time intervals (~56, 31, and 12 generations ago) using the MAPS software (Al-Asadi et al. 2019). The migration surfaces were correlated across the time periods (Pearson R=0.37-0.84) and across three independent runs (Pearson R=0.54-0.97) (Fig S1). The migration surfaces showed barriers to migration in southwestern Ethiopia and parts of Kenya and Tanzania (Fig. 3). The inferred migration rates between demes were relatively low, with most values ranging from ~0.001-10, corresponding to an estimated dispersal distance (distance traveled by an individual after one generation) of ~1km-400km. The migration surfaces were similar across the three time periods, although the most recent one, corresponding to 12.5 generations ago, showed higher migration around South Sudan and lower migration in southwestern Kenya and northern Tanzania compared to the older two. The surfaces also show that the model has some bias toward inferring lower migration in areas where genomic data were collected, perhaps because many of these populations were targeted for their unique language or subsistence strategy, especially in Ethiopia and Tanzania. A network depicting the sharing of identical by descent tracts recapitulates the complex patterns of connection shown in the MAPS output (Fig. S7).

**Figure 2:**
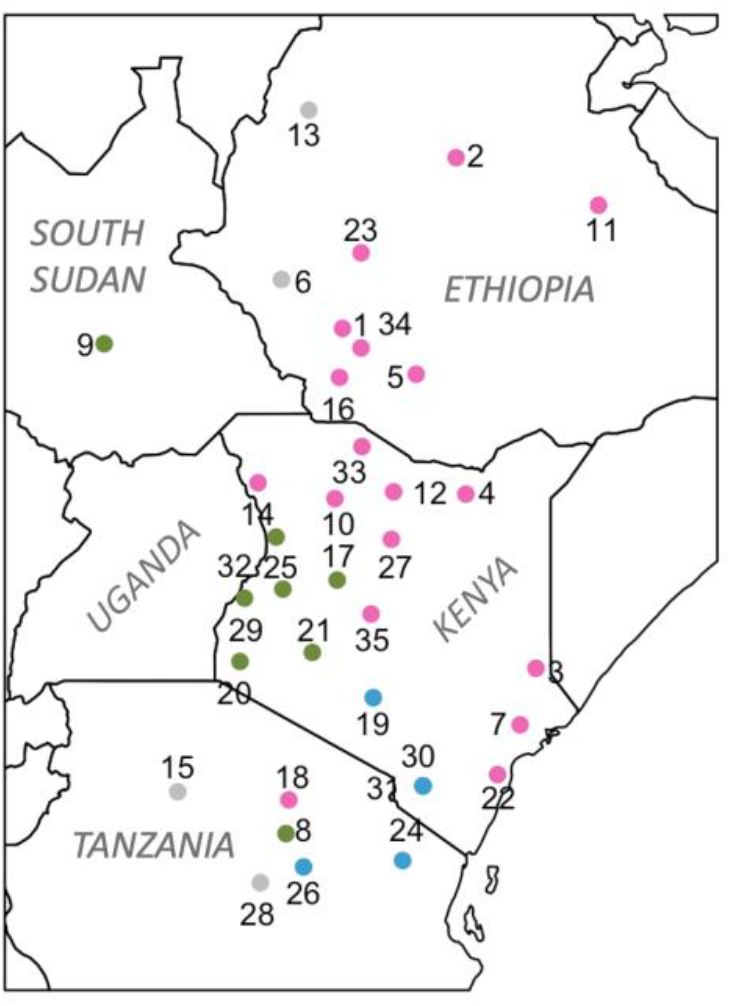
Geographic locations of genetic samples within eastern Africa included in the SPRUCE analysis (35 population locations, 492 individuals). Colors correspond to language family of the population: pink = Afro-Asiatic, blue = Niger-Congo, green = Nilo-Saharan, grey = isolate (6. Chabu, 13. Gumuz) or Khoisan (15. Hadza, 28. Sandawe). Numbers correspond to Table S1.

**Figure 3:**
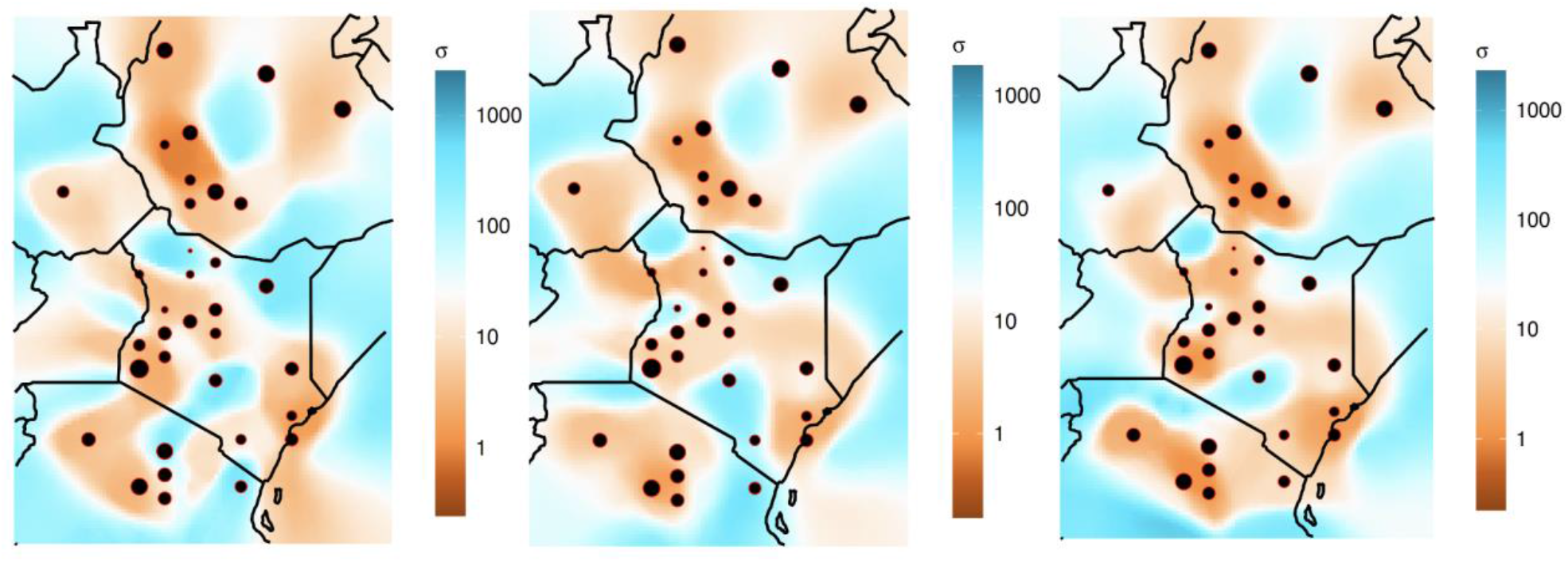
Inferred dispersal surfaces based on shared identical by descent tracts using the program MAPS (Al-Asadi et al. 2019). Time intervals correspond to ~56 generations ago (left), ~31 generations ago (middle), and ~12 generations ago (right). Light blue shows higher migration and brown shows lower migration. Estimates of migration were transformed into dispersal distance (σ) by scaling with the grid step-size (Al-Asadi et al. 2019).

We used a random forest framework to test the effects of environment, language, and an important disease vector (tsetse fly) presence on migration. Specifically, we extracted the inferred migration rate value (our response variable) from 368 evenly spaced locations in our region of interest along with the values at each of the same coordinates for each of the spatial variables (our predictor variables). Our random forest models explained 42.3%, 38.5%, and 46.6% of the variance for each of the three time periods from oldest to most recent. The Pearson R correlation between predicted and actual migration was >0.62 for all three models (Fig. 4). The most important variables across the three time periods were similar; minimum temperature of the coldest month and the kernel density surface were in the top four variables across the three models (Table 1). (The kernel density shows the geographic density of genetic samples and helps account for spatial autocorrelation in the model). Other important variables included mean annual precipitation, minimum precipitation of the driest month, maximum temperature of the hottest month, altitude, and presence of the Niger-Congo languages (see Figs. S2-S4 and Table S2 for more information on the predictor variables). To better understand the influence of the kernel density surface in the random forest model, we tried rerunning the model after excluding it. The model performance decreased slightly; the percent variance explained was >33% and the correlation between predicted and actual migration was >0.57 across the three time periods. The most important variables were similar to those from the full model, although the presence of the Niger-Congo languages increased in importance. Full results are available in Table S4.

**Figure 4:**
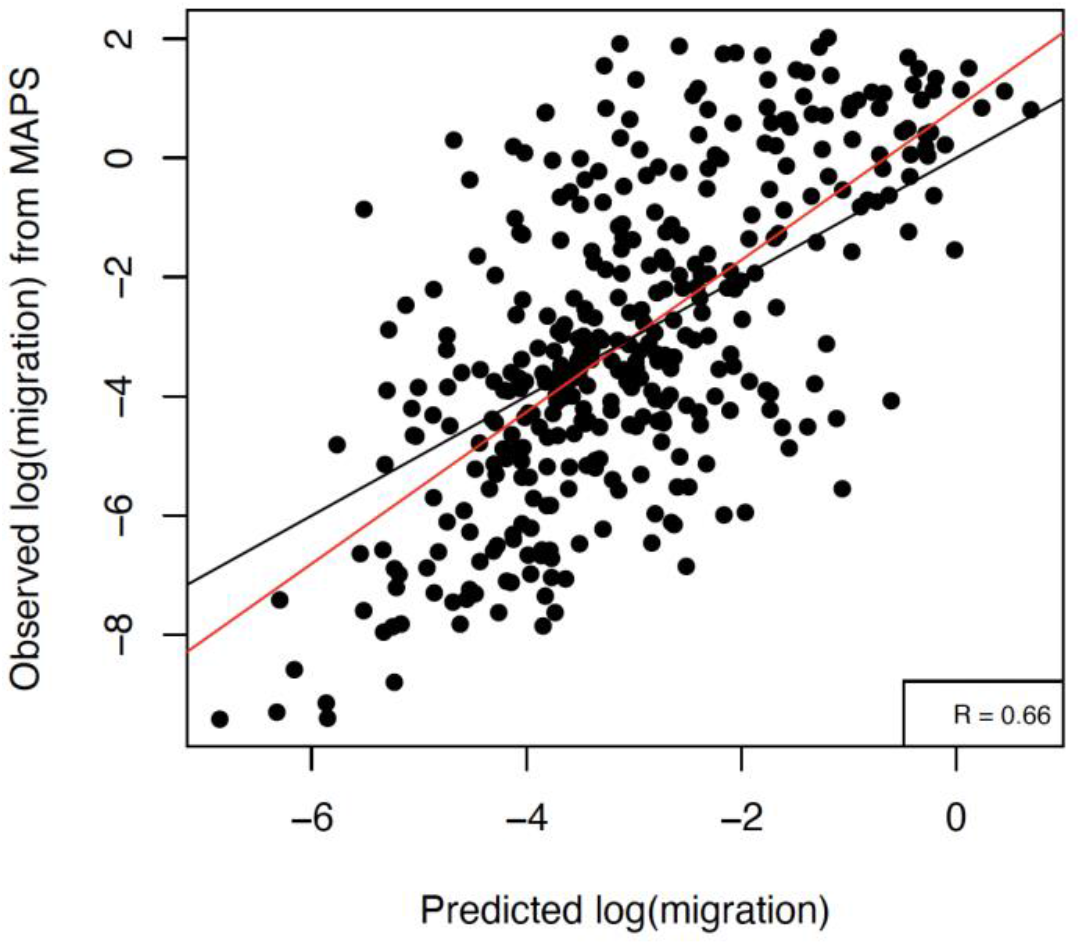
Observed migration (inferred by MAPS) versus predicted migration rate by SPRUCE. The random forest regression was trained on the full dataset for the oldest time period (~56 generations ago). The red line is the best-fit linear regression, and the black line is y=x.

**Table 1:** Summary of results from full dataset run across the three time intervals, with and without the kernel density surface included as a predictor variable. R = correlation between observed and predicted migration.

| Spatial variables | Time interval | Percent variance explained | R | Most important variables |
| --- | --- | --- | --- | --- |
| All | 2-4 | 42.3 | 0.664 | Kernel, MinTemp, MeanPrec, Altitude |
| All | 4-6 | 38.5 | 0.626 | Kernel, MinTemp, MeanPrec, PrecDry |
| All | 6-Inf | 46.6 | 0.682 | Kernel, LangNC, MinTemp, MaxTemp |
| Excluded kernel | 2-4 | 39.3 | 0.635 | MinTemp, MeanPrec, LangNC, Altitude |
| Excluded kernel | 4-6 | 33.3 | 0.591 | PrecDry, Altitude, MinTemp, LangNC |
| Excluded kernel | 6-Inf | 33.1 | 0.578 | PrecDry, LangAA, MinTemp, LangNC |

To ensure the model was not overfitting the data, we ran the random forest regression with a training dataset (70% of the data) and validated it with the testing dataset (30%) for all three time periods and with and without the kernel density variable. In the majority of runs, the correlation between predicted and actual migration was the same or even somewhat lower for the training dataset (R_train_) than for the test dataset (R_test_), indicating the model was not overfit (Table S5, Fig S5). The only exception was the model corresponding to the youngest period (12.5 generations ago) and built without the kernel density surface, in which case Rtrain (0.57) was higher than R_test_ (0.38).

### Selection Scan

Since the results from SPRUCE indicated that fine-scale environments may affect migration and selection, we decided to test this further by performing a genome-wide selection scan on several populations in southwestern Ethiopia who live close within 70km but derive from different ancestries and experience different environments. Four populations are agriculturalists, ranging from small-scale (Majang) to intensive farming (Shekkacho) (Fentaw 2007; Stauder 2007; Gopalan et al. 2022). One population, the Chabu, practiced traditional hunting and gathering until recently (Dira and Hewlett 2018; Gopalan et al. 2022). Some individuals in these populations live at moderately high altitudes; for example, the Chabu live at elevations up to 2500m (Dira and Hewlett 2017). For the selection scan we focused on the integrated Haplotype Score (iHS), a statistic which detects recent positive selection through decay of ancestral and derived alleles from a query locus (Voight et al. 2006). After filtering for the 0.1% largest absolute value iHS scores, we intersected these SNPs with a list of known genes, resulting in a range of 129 (Majang) to 230 (Shekkacho) top candidate genes (Dataset S1). Out of 664 unique genes found across the five populations, only six genes were candidate genes for all five populations. These six genes were related to metabolism (PTPRN2, RPH3AL), the immune system (DOCK8), cAMP binding (PRKAR1B), and vesicle-mediated transport (VPS53). We found five genes that were candidates for selection in the four agricultural populations but not in the hunter gatherer population (the Chabu); one with a possible relationship to subsistence strategy is PASK, which regulates insulin gene expression and can play a role in type 2 diabetes (Zhang et al. 2015). We also found that the agricultural groups shared a higher percentage of top candidate genes with each other than they did with the Chabu, with one exception (Majang was more similar to Chabu than to Shekkacho, reflecting their shared genetic ancestry) (Fig. 5, Table S7). The Chabu and Majang also shared a common candidate gene called IRF4, which is an important regulatory transcription factor in the development of immune cells. We did not find evidence of selection in genes previously associated with high altitude adaptation (EPAS1, ALPP, PTGIS, EGLN1, KCTD12, NOS2, and VDR). Interestingly, a subunit of the hypoxia-inducible factor 1 pathway (HIF1A) was detected as a candidate gene in the Chabu.

**Fig. 5:**
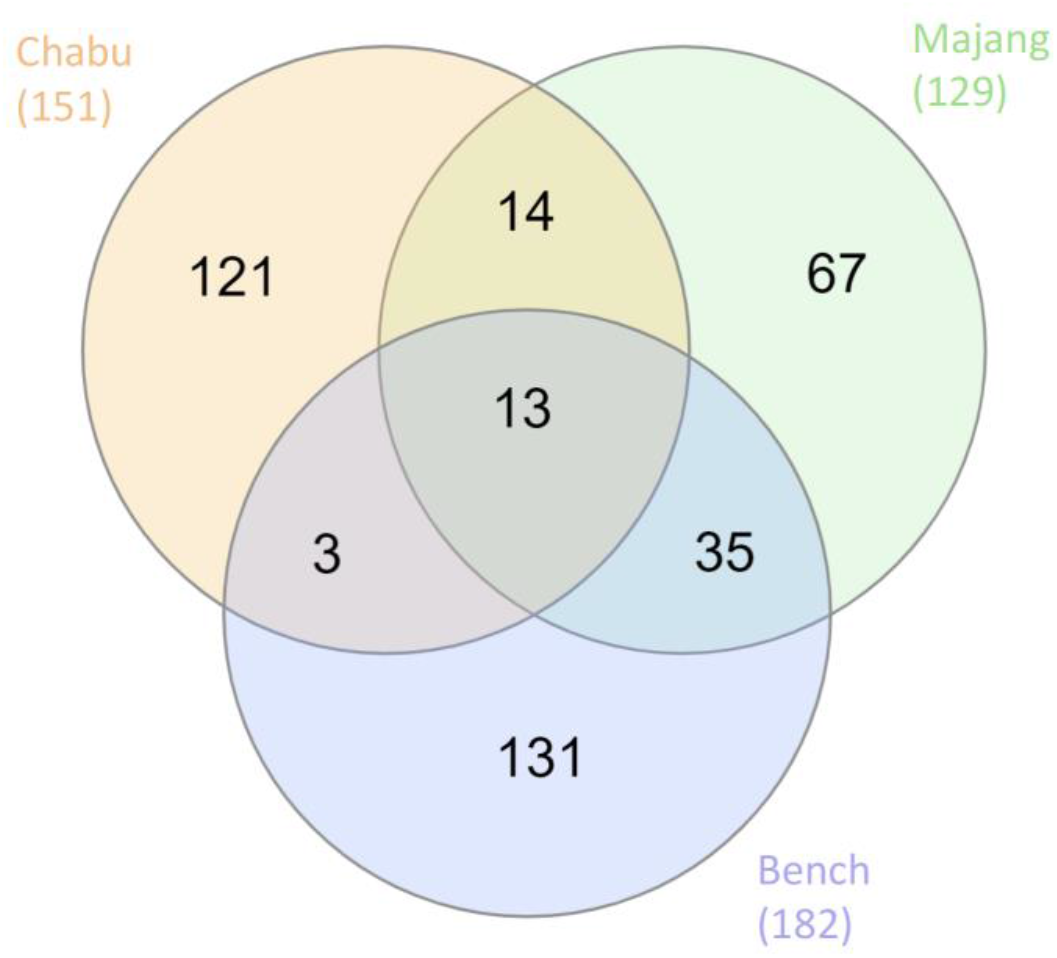
Similarity in selection scan results across three populations from southwestern Ethiopia. This Venn Diagram shows how many candidate genes are unique and shared among three populations (the Chabu are recent hunter-gatherers while the Bench and Majang are agriculturalists). Candidate genes were found using the largest 0.01% iHS scores (absolute value), and the total number of unique candidate genes found for each population is listed under the population name.

## Discussion

Here we present a new methodology for predicting the environmental drivers of human migration and gene flow. This method can be applied across a wide range of regions and scales and is also suitable for other organisms, given they have a reference genome and high-resolution genetic data suitable for calling identical by descent (IBD) tracts. We based our model on a previous method (Pless et al. 2021), which used an iterative random forest approach to infer genetic connectivity based on pairwise summary statistics (e.g. F_ST_) and the corresponding mean of environmental values between each of these pairs of populations. Our new method SPRUCE avoids the need for pairwise statistics by inferring migration rate directly through geographic space using the software MAPS (Al-Asadi et al. 2019). We applied our new method to investigate patterns of migration in eastern Africa 300-2,000 years ago and focused on different time periods by filtering for different length IBD tracts. We found a high degree of correlation across the time periods, which indicates that the evolutionary forces that caused IBD sharing in one time period overlap with those that affected IBD sharing in proximate time periods (Al-Asadi et al. 2019). In some regions, we found that low migration rates corresponded to geographic areas expected to be barriers, such as the mountainous forests of southwestern Ethiopian and the arid Lake Turkana region in northern Kenya. On the other hand, some regions that we expected to be barriers (the Indian Ocean coastline, the Ethiopian central highlands, Mount Kenya, and Mount Kilimanjaro) did not appear as barriers in the MAPS output. The lush and ecologically diverse Lake Victoria region did show evidence of promoting migration, and we observed a complex pattern of barriers and corridors in Kenya and Tanzania. For example, southeastern Kenya and northern Tanzania see strong migration between Iraqw and Kikuyu during the oldest period potentially reflecting the incursion of the Bantu-speaking agriculturalists into the region. Then, during the intermediate ~900 year period, we see a switch to higher migration among the Kikuyu, Taita and Pare (all of whom are Bantu-speakers). Finally, most recently, there is increasing isolation among these groups.

The 25 spatial variables representing the environment (climate, elevation, land cover), tsetse fly occurrence, and language family explained ~40% of the variance in migration rate and achieved an R of >0.62. Although this is lower than the R from our original model applied to a species of mosquito, it is on par with other landscape genetics papers that use machine learning (Murphy et al. 2010; Hether and Hoffman 2012; Fountain-Jones et al. 2017) and seems noteworthy given the immense complexity of human migration and the environment in eastern Africa. Minimum temperature, mean precipitation, and altitude were consistently three of the most important variables. In some regions, these variables were correlated with each other: high elevation regions tended to have higher precipitation and lower minimum temperatures. These conditions likely acted as a barrier to gene flow within southwestern Ethiopia and parts of southern Kenya, given low rates of IBD-sharing and inferred migration in these areas (Fig. 3, Fig. S7). Many of the southern Ethiopian populations (e.g. Chabu, Aari, Hamer, Oromo, Burji and Wolayta) cross-cut language families and subsistence strategies (Assoma 2007) and have extremely low migration rates (Fig. S7). In the most recent time period (12 generations ago), the presence/absence of the Niger-Congo languages was the second most important variable. This is likely driven by high migration areas along the southern and eastern border of our study region, which overlap with Bantu-speaking populations (Fig. S3), implying sharing this language family may promote intermarriage and migration among populations.

Another important variable in predicting migration rates was the kernel density surface, which has higher values in areas with higher sampling density (i.e. more individuals with genetic data). We included this variable to help account for spatial autocorrelation, which arose because our genetic samples were not uniformly distributed across the region. The kernel density surface was probably important because MAPS tended to predict low migration rates near genetic samples. This may be due to the presence of relatively isolated neighboring populations, while the high migration rates around the sampling locations were necessary to explain instances of IBD sharing across large geographic regions (Fig. S7). Interestingly, we noticed a similar bias when investigating migration barriers using the software that preceded MAPS (EEMS, (Petkova et al. 2016)) for southern Africa, an area that also has complex migration and isolated populations (Petkova et al. 2016; Uren et al. 2016). When we excluded the kernel density surface from our model, precipitation of the driest month and Niger-Congo language presence/absence rose in importance.

In the case of the intermediate time-period model, corresponding to 31 generations ago, one of the most important variables was the occurrence of a tsetse fly group called fusca (forest-dwelling), and the second most import tsetse fly group was morsitans (savannah-dwelling) (Fig. S4). The tsetse fly is the vector of livestock trypanosomiasis and sleeping sickness in humans, and its occurrence likely limits the movement and health of pastoralist groups (Gifford-Gonzalez 2017). Cattle herding spread into Lake Turkana (northern Kenya) from Sudan by 5,000-4,000 ya (Hildebrand and Grillo 2012) and remains a frequent subsistence strategy among many present-day pastoralists. The Kenyan and Sudanese cattle pastoralist populations occupy areas without morsitans (Fig. S4a), and many of these groups share high cross-population IBD such as the Gabra, Borana, Rendille and Gurreh (Fig. S7). The fusca tsetse fly range is heavily represented in forested areas of southwestern Ethiopia. This may have served as a barrier to gene flow preventing Sudanese and northern Kenyan pastoralists from major incursions into the highlands.

Since the SPRUCE results indicated that fine-scale differences in environment affect human migration, we ran a genome-wide selection scan on five neighboring populations in southwestern Ethiopia to test the effect of microenvironments on adaptation. Although these populations are close in geographic space, and in some cases closely related (Gopalan et al. 2022), they each show a unique pattern of positive selection using the iHS statistic (Fig. S6). We predicted that differences in pathogen exposure, subsistence strategies, and elevation would be the primary drivers of differential selection in this region, and indeed a number of the candidate genes were related to the immune response (e.g. IRF4, AXIN1, CHL1, DOCK8) or metabolism and insulin production (e.g. PNPLA2, PASK, RPH3AL, B4GALNT3). Additionally, the pairs of the agricultural groups shared more candidate genes in common than almost any of them do with the recent hunter-gatherer group (Table S7). We did not find evidence of selection for any of the specific genes previously associated with high altitude adaptation in the literature. Two possible explanations are 1) less selection pressure due to lower elevations in Ethiopia than the Andes or Tibetan plateau, and 2) selection scans do not replicate well in African populations (Granka et al. 2012). We do highlight one candidate gene in the Chabu called HIF1A which is a part the hypoxia-inducible factor 1 pathway. Several other HIF genes have been identified as candidate genes for adaptation to high altitude, in particular EGLN1, an O_2_ sensor which controls levels of HIF-alpha (Bigham et al. 2013; Brutsaert et al. 2019).

Like all models, SPRUCE has certain limitations and will benefit from additional work. One of the limitations of this method is that it relies strongly on the output of another software program to predict migration rates. A pattern of complex relationships across many geographically separated individuals is inherently difficult to represent through one two-dimensional migration surface, and the migration surfaces in this study are certainly complicated and serpentine in appearance. Another option we considered is to use IBD sharing as the predictor variable in the model, although this adds the complexity and inconvenience of using pairwise summaries of environmental variables, rather than using values corresponding to a single location. Another option is to consider other statistical frameworks that may improve the model’s interpretability. However, random forest has the advantage that it allows for a high level of objectivity since it can accept many predictor variables, included correlated variables and both continuous and categorical variables (Breiman 2001; Hastie et al. 2009).

The flexibility of SPRUCE can help users overcome some of its limitations. Not only could the user replace random forest with another statistical framework, but they could experiment with other tools to generate the response variable (migration rates). For the predictor variables (e.g. environmental data), there are a wide variety of open-source spatial rasters, or the user could generate their own spatial variables of interest including distance to a hypothesized corridor for gene flow (e.g. the coast or a river). Another improvement would be to generate spatial variables that correspond directly to the time period under investigation, for example using historical climate reconstruction to create the climate rasters (Singarayer and Valdes 2010; Beyer et al. 2021). A current challenge to this is that historical climate reconstructions are generally performed at a coarse time-scale, e.g. in 1,000 year intervals. Given the relatively short time scale we consider in this project, we think it is reasonable to assume that the variables we included have not changed drastically in the last 2,000 years, although there have been some important changes such as a slight cooling ~1,500 years ago followed by warming ~1,100 years ago (Nicholson et al. 2013). Additionally, applying SPRUCE to study human migration in other regions would be an interesting way to examine which environmental variables are broadly relevant to human migration, and which are “limiting factors” depending on the ecological and temporal context.

In conclusion, we find evidence of fine-scale effects of environment on human migration and adaptation in eastern Africa, and we present a new approach for integrating genetic and environmental data to predict the ecological drivers of population structure and migration.

## Materials and Methods

### SPRUCE Model

We provide the full scripting procedure and additional commentary on SPRUCE here: https://github.com/evlynpless/SPRUCE-RF. MAPS software and resources can be found here: https://github.com/halasadi/MAPS.

#### Environmental and spatial data

Climate and landcover data were downloaded from free, online repositories and were edited and cropped using Geospatial Data Abstraction Library (GDAL/OGR contributors 2021) under the Bash environment. Most datasets were available at 1-km resolution; otherwise, they were resampled to a pixel size of 1 km^2^. Elevation was derived from MERIT DEM (Multi-Error-Removed Improved Terrain Digital Elevation Models) (Yamazaki et al. 2017), and slope was derived from the Geomorpho90m dataset (Amatulli et al. 2020). Mean annual temperature, maximum temperature of the hottest month, minimum temperature of the coldest month, mean precipitation of the wettest month, and mean precipitation of the driest month were obtained from CHELSA (climatologies at high resolution for the earth’s land surface areas) (Karger et al. 2017). We also included aridity from the Global Aridity Index, and gross primary production, a measure of vegetative photosynthesis (Zomer et al. 2007; Zomer et al. 2008). Landcover types are from one source (Tuanmu and Jetz 2014) but each represented by a separate raster, in which values from 0 to 1 indicate what percent of that pixel is covered with each landcover type.

Tsetse suitability maps for the three main group (morsitans, fusca, and palpalis) were download as shapefiles from the Food and Agriculture Organization of the United Nations (Cecchi et al. 2008) and converted to rasters using qGIS 2.18 (QGIS Development Team 2017). Language data (including language family and geographic coordinates) were downloaded from Ethiolang and Wikitongues and categorized by language family using R Statistical Software (v.4.0.2; R Core Team 2020). Local convex hulls were created for the three primary language families in eastern Africa, Afro-Asiatic, Nilo-Saharan, and Niger-Congo, using the R package “LoCoH” (Fig. S3) (Getz et al. 2007). To address spatial autocorrelation among the geographic coordinates of the individuals, we included a kernel density raster (bandwidth 200 km) created using the R package “KernSmooth” (Wand et al. 2015).

The values at each deme for each spatial variable were extracted with the R package rgdal (“gdalinlocationinfo”) (Bivand et al. 2021), with the exception of the language maps which were extracted using the R package sf (“st_contains”) (Pebesma 2018). The location and correct order of the demes to match the inferred migration rates was found in the demes.txt output file from the MAPS software.

#### Genomic data, phasing, and IBD

For the SPRUCE analysis, we included dense SNP array data from 35 populations in eastern Africa from two sources (Gurdasani et al. 2015; Scheinfeldt et al. 2019) (Table S1). These populations were primarily from Ethiopia, Kenya, and Tanzania and include ten populations that practice or recently practiced hunting and gathering as their primary subsistence strategy. In preparation for merging the two datasets, all variants were oriented to match the 1000 Genomes reference using a custom script. We flipped SNPs that were on the wrong strand, and we excluded SNPs that did not match the 1000 Genomes reference. We merged the datasets using plink1.9 (--merge-mode 1) and removed loci with genotype missingness >95% (Purcell et al. 2007; Purcell 2016). The merged dataset included 517,383 SNPs and 821 individuals from 50 populations. We performed principal component analysis (PCA) with plink, which confirmed the relationships among the populations were as expected (Fig. S6). In preparation for calling identical by descent (IBD) segments, we performed phasing with SHAPEIT2 using a reference panel of phased individuals from the 1000 Genomes project Phase 3 dataset, the --duohmm option, and a window size of 5 Mb (Delaneau et al. 2013). We then removed first and second degree relatives identified by KING (Manichaikul et al. 2010). Shared IBD segments were obtained with hap-ibd v1.0 (Zhou et al. 2020) and repaired with the program merge-ibd-segments. We excluded 15 populations that fell outside our geographic region of interest (top left = 12.5°N, 30.0°E, bottom right= −7.0°S, 44.0°E), leaving 35 populations and 492 individuals for the SPRUCE analysis (Fig. 2).

To visualize patterns of IBD sharing, we created networks in Cytoscape 3.9.1. (Shannon et al. 2003). In these networks, each individual is a node, and the locations of the nodes roughly correspond to geographic positions. The edges represent number of shared IBD segments, and for ease of interpretation, we focused on IBD segments >6cM (corresponding to the most recent time interval considered in MAPS).

#### Population names and coordinates

Throughout the manuscript we use population names from the original sources with the exception of the “Sabue” which we refer to as “Chabu” (Dira and Hewlett 2017). Likewise, we use geographic coordinates from the original publications when available. Coordinates that were not included in Gurdasani et. al (2015) (or the original source, Pagani et al. 2012)) were taken from Gopalan et al. (2021). Coordinates not included in Scheinfeldt et al. (2019) were inferred from a map in the manuscript using a custom script.

#### MAPS

In preparation for running MAPS, we adapted and ran code provided by the program authors to convert the output of hap-ibd into a matrix format read by MAPS, and in the process we filtered IBD calls into three categories: 2-4cM, 4-6cM, and >6cM. Under a simplistic model of infinite population size, the mean ages of the categories correspond to 56.25, 31.25, and 12.5 generations respectively (Al-Asadi et al. 2019). We ran MAPS with 400 demes, and we averaged inferred migration over three replicates to mediate any outlier results. The number of MCMC iterations in each replicate was set to 5 million, the burn-in was set to 2 million, and we thinned every 2,000 iterations as in Al-Asadi et al. (2019). Migration surfaces were visualized using the R package “plotmaps” (Al-Asadi et al. 2017). For easier interpretation, we also report dispersal distance, which is the effective spatial diffusion parameter, calculated by scaling migration by the step size of the grid used by the MAPS software (Al-Asadi et al. 2019). Dispersal distance is roughly equal to the expected distance an individual disperses in one generation (Al-Asadi et al. 2019).

#### SPRUCE

The SPRUCE analysis used a random forest regression performed with the R package randomForestSRC (Ishwaran et al. 2022). We tuned the random forest regression for the number features to consider at each split point (“mtry”) and the minimum size of terminal nodes (“nodesize”). We calculated a migration rate at each deme by taking values in the “mrates.txt” output file and raising them to the tenth power (Al-Asadi, personal communication). Since migration rates were unbalanced among populations and skewed toward low values, we used a log transformation before including them in the model (Fig. S8). The predictor variables were the values of 25 spatial variables related to climate, ecology, disease vectors, and language at each deme, as well as the kernel density surface (Table S2). All variables were numeric and continuous except for language which was coded as “1” for language family present, and “0” for language family not present. We performed random forest regressions for each of the three time intervals of interest (56.25, 31.25, and 12.5 generations ago). We ran the analyses with and without including the kernel density surface as a predictor variable to better understand its influence on the model. Finally, we validated the model by withholding a test dataset (30%) and rerunning all regressions.

### Genome-wide Selection Scans

#### Genomic data

While the SPRUCE dataset described above gave us the broadest possible geographic spread in eastern Africa, some populations had relatively low sample sizes. We therefore considered an alternative high-SNP density, high sample size dataset of five Ethiopian populations previously published in (Gopalan et al. 2022) and available in dbGaP at phs001123.v2.p2. We tested for selection in these five populations from southwestern Ethiopia: Bench, Sheko, Majang, Shekkacho, and Chabu. Fifty individuals from each population (and 88 from the Chabu) were genotyped on the Illumina Infinium MultiEthnic Global Array, which assays over 1.7 million genetic markers. Gopalan et al. filtered SNPs to remove samples with call rates <90%. Variants with MAF <5% were updated with zCall genotypes, and then variants were removed if they had >15% missing data, heterozygosity >= to 80%, or cluster separation <= to 2%.

#### Selection scan

To prepare genetic data for haplotype-based selection scans, we implemented phasing using a reference panel of phased individuals from the 1000 Genomes project Phase 3 dataset (The 1000 Genomes Consortium 2015), the --duohmm option, and a window size of 5 Mb in SHAPEIT2 (Delaneau et al. 2013). PONDEROSA (Williams et al. 2020), an algorithm for accurately recovering family-level relationships from genome-wide data, was used to exclude individuals with a second degree or closer relative in the dataset, leaving 222 individuals. We calculated integrated Haplotype Score (iHS) (Voight et al. 2006), using selscan 1.3.0 (Szpiech and Hernandez 2014) and default parameters and a genetic map based on crossovers in 30,000 unrelated African Americans (Hinch et al. 2011). We found the absolute value of the normalized iHS scores and filtered for the 0.1% largest absolute value iHS scores. We annotated these genetic positions with a gene range list provided by Plink (https://www.cog-genomics.org/plink/1.9/resources).

To compare top gene hits across the five populations, we created Venn Diagrams using InteractiVenn (Heberle et al. 2015). We generated plots showing iHS scores across the genome using the R package “rehh” (Gautier and Vitalis 2012).

## Supporting information

Supplemental Informaion

## Acknowledgements

We thank all communities that have contributed data to the studies included in this analysis, and we thank Dr. Luca Pagani and Dr. Sarah Tishkoff for making their datasets available to us. We acknowledge the sovereignty and rights of all of these groups to the governance, protection, and use of their own genetic data. We thank Dr. John Novembre and Dr. Al-Asadi for providing consultation on the MAPS software program. We thank Dr. Shyamalika Gopalan for advice and support throughout the project, Dr. Richard Berl for his help in generating the language maps, and Dr. Cesar Augusto Fortes-Lima and Dr. Carina Schlebusch for their help with population metadata. We also thank Dr. Ackmel Negash, Justin Myrick, and Mira Mastoras for helpful discussions and the UC Davis Data Lab and R User Group for advice on statistics and coding. This research was supported by NIH grant R35GM133531 (to BMH). The content is solely the responsibility of the authors and does not necessarily represent the official views of the National Institutes of Health.

## Notes

### Competing Interest Statement

The authors have declared no competing interest.

### Summary of Updates

Updated and corrected references

