## Supplemental Informaion for "Predicting environmental and ecological drivers of human population structure"

[1]Department of Anthropology, University of California, Davis
[2]UC Davis Genome Center, University of California, Davis
*Corresponding authors

**The supplementary materials include:**

Tables S1 to S7
Figures S1 to S8
Legend for Dataset S1
SI References

**Other supplementary materials for this manuscript:**

Dataset S1

**Table S1.** Populations included in SPRUCE model. Number (consistent with Fig. 1), population name, country, language family, traditional subsistence strategy, longitude, latitude, number of individuals, and reference: [1] is Scheinfeldt et al. 2019, [2] is Gurdasani et al. 2015 (originally collected for Pagani et al. 2012).

| Number | Population | Country | Language | Subsistence | Longitude | Latitude | N | Reference |
|---|---|---|---|---|---|---|---|---|
| 1 | Aari | Ethiopia | Afro-Asiatic | Agropastoralist | 36.6106 | 6.4108 | 10 | [1] |
| 2 | Amhara | Ethiopia | Afro-Asiatic | Agriculturalist and/or pastoralist | 39 | 10 | 25 | [2] |
| 3 | Boni | Kenya | Afro-Asiatic | Hunter-Gatherer | 40.6811 | -0.7333 | 15 | [1] |
| 4 | Borana | Kenya | Afro-Asiatic | Pastoralist | 39.2069 | 2.9284 | 18 | [1] |
| 5 | Burji | Kenya | Afro-Asiatic | Agriculturalist | 38.1608 | 5.4487 | 14 | [1] |
| 6 | Chabu | Ethiopia | Nilo-Saharan / Isolate | Hunter-Gatherer | 35.3239 | 7.4402 | 8 | [1] |
| 7 | Dahalo | Kenya | Afro-Asiatic | Hunter-Gatherer | 40.3482 | -1.9221 | 9 | [1] |
| 8 | Datog | Tanzania | Nilo-Saharan | Pastoralist | 35.42 | -4.21 | 16 | [1] |
| 9 | Dinka | Sudan | Nilo-Saharan | Pastoralist | 31.5924 | 6.0891 | 12 | [1] |
| 10 | Elmolo | Kenya | Afro-Asiatic | Hunter-Gatherer | 36.4488 | 2.8333 | 6 | [1] |
| 11 | Ethiopian Somali | Ethiopia | Afro-Asiatic | Agriculturalist and/or pastoralist | 42 | 9 | 24 | [2] |
| 12 | Gabra | Kenya | Afro-Asiatic | Pastoralist | 37.6852 | 2.9759 | 9 | [1] |
| 13 | Gumuz | Ethiopia | Isolate | Horticulturalist | 35.9 | 11 | 22 | [2] |
| 14 | Gurreh | Kenya | Afro-Asiatic | Pastoralist | 34.832 | 3.1661 | 7 | [1] |
| 15 | Hadza | Tanzania | Khoisan | Hunter-Gatherer | 33.14 | -3.33 | 16 | [1] |
| 16 | Hamer | Ethiopia | Afro-Asiatic | Pastoralist | 36.5463 | 5.3814 | 10 | [1] |
| 17 | Ilchamus | Kenya | Nilo-Saharan | Pastoralist | 36.4964 | 1.1213 | 16 | [1] |
| 18 | Iraqw | Tanzania | Afro-Asiatic | Agriculturalist | 35.48 | -3.5 | 22 | [1] |
| 19 | Kikuyu | Kenya | Niger-Kordofanian | Agriculturalist | 37.2572 | -1.3515 | 14 | [1] |
| 20 | Luo | Kenya | Nilo-Saharan | Pastoralist | 34.4516 | -0.5906 | 31 | [1] |
| 21 | Ogiek | Kenya | Nilo-Saharan | Hunter-Gatherer | 35.9733 | -0.4004 | 12 | [1] |
| 22 | Orma | Kenya | Afro-Asiatic | Pastoralist | 39.8727 | -2.9683 | 14 | [1] |
| 23 | Oromo | Ethiopia | Afro-Asiatic | Agropastoralism | 37 | 8 | 20 | [2] |
| 24 | Pare | Kenya | Niger-Kordofanian | Agriculturalist | 37.8754 | -4.7753 | 12 | [1] |
| 25 | Pokot | Kenya | Nilo-Saharan | Pastoralist | 35.2124 | 2.0249 | 5 | [1] |
| 26 | Rangi | Tanzania | Niger-Kordofanian | Agriculturalist | 35.79 | -4.91 | 14 | [1] |
| 27 | Rendille | Kenya | Afro-Asiatic | Pastoralist | 37.6377 | 1.9773 | 14 | [1] |
| 28 | Sandawe | Tanzania | Khoisan | Hunter-Gatherer | 34.88 | -5.24 | 24 | [1] |
| 29 | Sengwer | Kenya | Nilo-Saharan | Hunter-Gatherer | 34.5467 | 0.7409 | 12 | [1] |
| 30 | Taita | Kenya | Niger-Kordofanian | Agropastoralist | 38.3034 | -3.2061 | 7 | [1] |
| 31 | Taveta | Kenya | Niger-Kordofanian | Agropastoralist | 38.3034 | -3.2061 | 2 | [1] |
| 32 | Tugen | Kenya | Nilo-Saharan | Pastoralist | 35.3551 | 0.9311 | 15 | [1] |
| 33 | Wata | Kenya | Afro-Asiatic | Hunter-Gatherer | 37.0195 | 3.927 | 4 | [1] |
| 34 | Wolayta | Ethiopia | Afro-Asiatic | Agropastoralist | 37 | 6 | 23 | [2] |
| 35 | Yaaku | Kenya | Afro-Asiatic | Hunter-Gatherer | 37.2097 | 0.408 | 10 | [1] |

**Table S2.** List of spatial variables included as predictor variables in the SPRUCE model. Abbreviation, description, source, resolution of the original dataset and resampling method used (if any).

| **Abbreviation** | **Description** | **Source name** | **Original resolution** |
|---|---|---|---|
| Aridity | Global Aridity Index | CGIAR CSI | $1km^2$ |
| MeanTemp | Annual mean temperature | CHELSA climate data | $1km^2$ |
| MaxTemp | Maximum temperature of the warmest month | CHELSA climate data | $1km^2$ |
| MinTemp | Coldest temperature of the coldest month | CHELSA climate data | $1km^2$ |
| MeanPrec | Annual precipitation | CHELSA climate data | $1km^2$ |
| PrecWet | Precipitation of the wettest month | CHELSA climate data | $1km^2$ |
| PrecDry | Precipitation of the driest month | CHELSA climate data | $1km^2$ |
| Evergreen | Evergreen/deciduous needleleaf trees (%) | Tuanmu and Jetz 2014 | $1km^2$ |
| Decid | Deciduous broadleaf trees (%) | Tuanmu and Jetz 2014 | $1km^2$ |
| Tree | Mixed/other trees (%) | Tuanmu and Jetz 2014 | $1km^2$ |
| Shrub | Shrubs (%) | Tuanmu and Jetz 2014 | $1km^2$ |
| Herb | Herbaceous vegetation (%) | Tuanmu and Jetz 2014 | $1km^2$ |
| Crop | Cultivated and managed vegetation (%) | Tuanmu and Jetz 2014 | $1km^2$ |
| Barren | Barren including spare shrub/herbaceous cover (%) | Tuanmu and Jetz 2014 | $1km^2$ |
| Water | Open water (%) | Tuanmu and Jetz 2014 | $1km^2$ |
| Slope | Slope | Amatulli *et al.* 2020 | $90m^2$, resampled by taking mean of pixels in $1km^2$ |
| Altitude | Altitude | MERIT DEM | $90m^2$, resampled by taking mean of pixels in $1km^2$ |
| GPP | Gross primary production, a measure of vegetation photosynthesis | Zhang *et al.* 2017 | $500m^2$, bilinear resampling |
| Fusc | Predicted area of suitability for tsetse flies (fusca): vector of livestock trypanosomiasis | Cecchi 2002 | 5km, not resampled |
| Mors | Predicted area of suitability for tsetse flies (morsitans): primary vector of T. rhodesiense in east Africa (mostly infect wildlife) | Cecchi 2002 | 5km, not resampled |
| Palp | Predicted area of suitability for tsetse flies (palpalis): primary vector of T. gambiense in west and central Africa | Cecchi 2002 | 5km, not resampled |
| LangAA | Inferred distribution of Afroasiastic languages | Glottolog and Wikitongues | NA (vector file) |
| LangNC | Inferred distribution of Niger-Congo languages | Glottolog and Wikitongues | NA (vector file) |
| LangNS | Inferred distribution of Nilo-Saharan languages | Glottolog and Wikitongues | NA (vector file) |
| Kernel | Kernel density map of sampled sites (bandwidth = 200km) | NA | $1km^2$ |

**Table S3**. Description and equations for each performance metric recorded for the model. The equations reference the randomforestSRC package in R and variables are defined as follows: RF = Random Forest model under consideration, M = migration rate (inferred by MAPS), TestingData = 70% of the dataset, ValidationData = 30% of the dataset.

| **Metrics for 10-fold cross-validation / training and validation models** | | |
|---|---|---|
| **Abbreviation** | **Description** | **Equation** |
| RSQ | Psuedo R-squared (percent variance explained by model, calculated by R package) | 1 - mse / Var(y) |
| $RMSE_{train}$ | Root mean square error of model for training dataset | sqrt(mean((RF$predicted.oob - TrainingData$M)^2)) |
| $RMSE_{valid}$ | Root mean square error of model for validation dataset | sqrt(mean((predict.rfsrc(RF, ValidationData)$predicted - TestingData$M)^2)) |
| $R_{train}$ | Pearson correlation between predicted and observed genetic distance for training dataset | cor(RF$predicted.oob,TrainingData$M) |
| $R_{valid}$ | Pearson correlation between predicted and observed genetic distance for validation dataset | cor((predict.rfsrc(RF, ValidationData))$predicted, ValidationData$M) |
| **Metrics for full dataset run** | | |
| **Abbreviation** | **Description** | **Equation** |
| $RSQ_{full}$ | Psuedo R-squared (percent variance explained by model, calculated by R package) | 1 - mse / Var(y) |
| $RMSE_{full}$ | Root mean square error of model | sqrt(mean((RF$predicted.oob - FullData$M)^2)) |
| $R_{full}$ | Pearson correlation between predicted and observed genetic distance for full dataset | cor(RF$predicted.oob, FullData$M) |

**Table S4.** Results for full dataset run. **RSQ** = R-squared (percent variance explained by the model); **$RMSE_{train}$** = root mean squared error of model for training dataset; **$RMSE_{valid}$** = root mean squared error of model for validation dataset; **$R_{train}$** = correlation between observed and predicted migration using training dataset; **$R_{valid}$** = correlation between observed and predicted migration using validation dataset; **Most important variables** are the four most important variables. For detailed information about these metrics, see Table S3.

| **Spatial variables** | **IBD cM** | **RSQ** | **RMSE** | **R** | **Most important variables** |
|---|---|---|---|---|---|
| All | 2-4 | 42.3 | 2.02 | 0.664 | Kernel, MinTemp, MeanPrec, Altitude |
| All | 4-6 | 38.5 | 2.18 | 0.626 | Kernel, MinTemp, MeanPrec, PrecDry |
| All | 6-Inf | 46.6 | 2.17 | 0.682 | Kernel, LangNC, MinTemp, MaxTemp |
| Excluded kernel | 2-4 | 39.3 | 2.07 | 0.635 | MinTemp, MeanPrec, LangNC, Altitude |
| Excluded kernel | 4-6 | 33.3 | 2.27 | 0.591 | PrecDry, Altitude, MinTemp, LangNC |
| Excluded kernel | 6-Inf | 33.1 | 2.43 | 0.578 | PrecDry, LangAA, MinTemp, LangNC |

**Table S5.** Results for training-validation run with all spatial variables included. **RSQ** = R-squared (percent variance explained by the model); **$RMSE_{train}$** = root mean squared error of model for training dataset; **$RMSE_{valid}$** = root mean squared error of model for validation dataset; **$R_{train}$** = correlation between observed and predicted migration using training dataset; **$R_{valid}$** = correlation between observed and predicted migration using validation dataset; **Most important variables** are the four most important variables. For detailed information about these metrics, see Table S3.

| **Spatial variables** | **IBD cM** | **RSQ** | **$RMSE_{train}$** | **$RMSE_{valid}$** | **$R_{train}$** | **$R_{valid}$** | **Most important variables** |
|---|---|---|---|---|---|---|---|
| All | 2-4 | 37.3 | 2.07 | 2.17 | 0.617 | 0.617 | Kernel, MinTemp, Altitude, PrecDry |
| All | 4-6 | 29.2 | 2.31 | 2.27 | 0.544 | 0.628 | Kernel, MinTemp, PrecDry, Decid |
| All | 6-Inf | 38.9 | 2.30 | 2.17 | 0.627 | 0.735 | Kernel, MinTemp, PrecDry, GPP |
| Excluded kernel | 2-4 | 35.6 | 2.13 | 2.10 | 0.610 | 0.634 | MinTemp, PrecDry, MeanTemp, Altitude |
| Excluded kernel | 4-6 | 21.35 | 2.47 | 2.28 | 0.462 | 0.591 | PrecDry, LangNC, MinTemp, Fusc |
| Excluded kernel | 6-Inf | 33.2 | 2.45 | 2.73 | 0.574 | 0.385 | PrecDry, LangAA, PrecWet, LangNS |

**Table S6.** Populations from southwestern Ethiopia included in the selection scan analysis. Population name, longitude, latitude, number of individuals, current or recent subsistence strategy, language family, and reference.

| Population | Longitude | Latitude | N | Subsistence | Language | Reference |
|---|---|---|---|---|---|---|
| Bench | 35.6 | 7 | 48 | Agriculture | Afro-Asiatic | Gopalan et al. 2021 |
| Chabu | 35.2 | 7.5 | 83 | Hunter-Gatherer | Nilo-Saharan / Isolate | Gopalan et al. 2021 |
| Majang | 35.3 | 7.2 | 49 | Agriculture | Nilo-Saharan | Gopalan et al. 2021 |
| Shekkacho | 35.4 | 7.2 | 46 | Agriculture | Afro-Asiatic | Gopalan et al. 2021 |
| Sheko | 35.5 | 7 | 50 | Agriculture | Afro-Asiatic | Gopalan et al. 2021 |

**Table S7**: Similarity in genome-wide selection scan results between Ethiopian populations. Specifically, this table shows the percentage of overlap in top candidate genes found using normalized integrated Haplotype Scores (iHS). The agricultural groups shared a higher percentage of top candidate genes with each other than they did with the hunter gather group (Chabu), with one exception shown in bold.

| | **Chabu** | **Bench** | **Majang** | **Sheko** | **Shekkacho** |
|---|---|---|---|---|---|
| **Chabu** | | 5% | 10.7% | 7.4% | 2.4% |
| **Bench** | 5% | | 18.3% | 21% | 8.1% |
| **Majang** | 10.7% | 18.3% | | 13.6% | **6.5%** |
| **Sheko** | 7.4% | 21% | 13.6% | | 8.4% |
| **Shekkacho** | 2.4% | 8.1% | **6.5%** | 8.4% | |

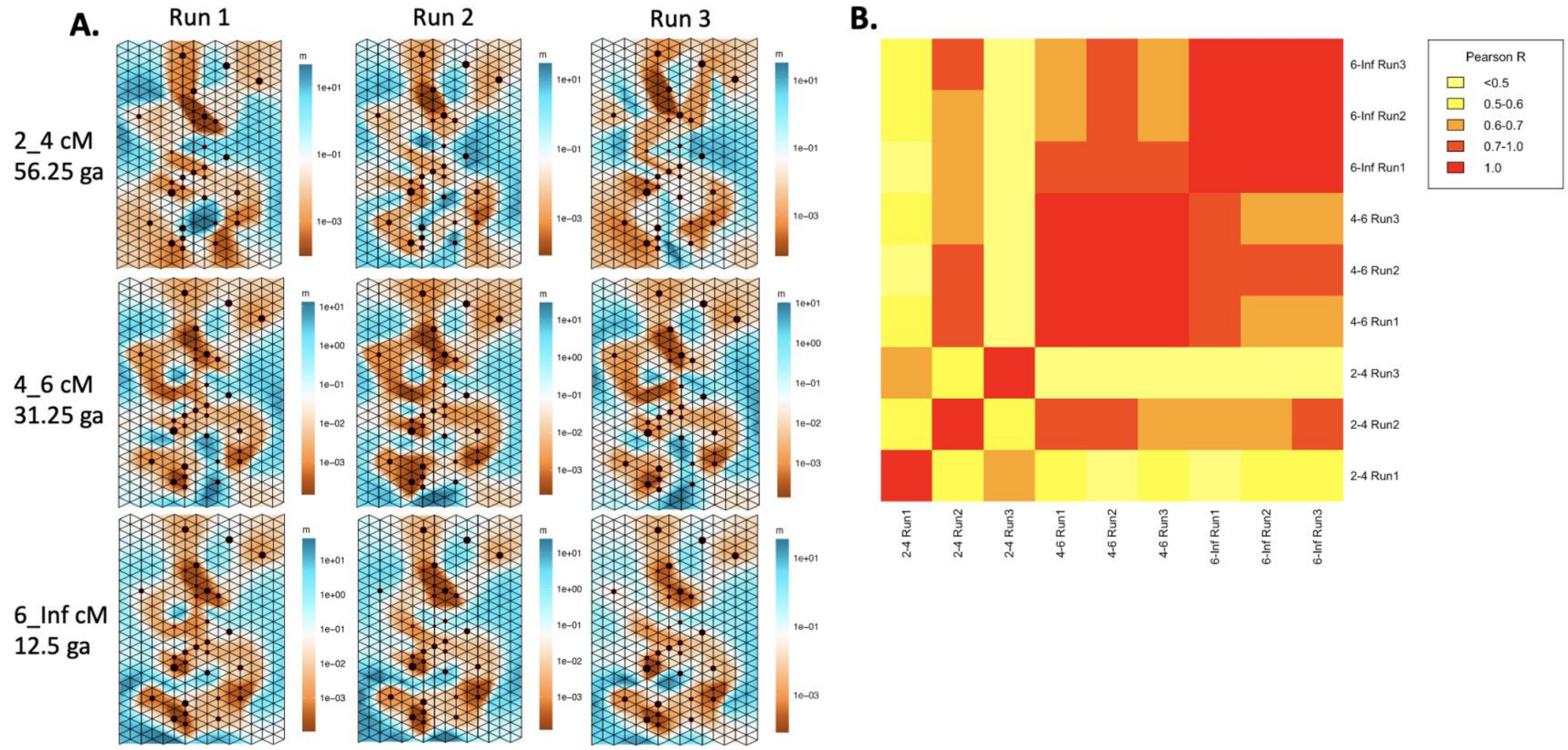


**Figure S1**: Migration surfaces generated by plotmaps (Al-Asadi et al. 2017), showing high correlation across runs and across time periods. Each column shows an independent run of the MAPS software (Al-Asadi et al. 2019), and each row represents a different time period in ga (generations ago) (i.e. different length identical by descent (IBD) tracts measured in cM (centimorgans)).

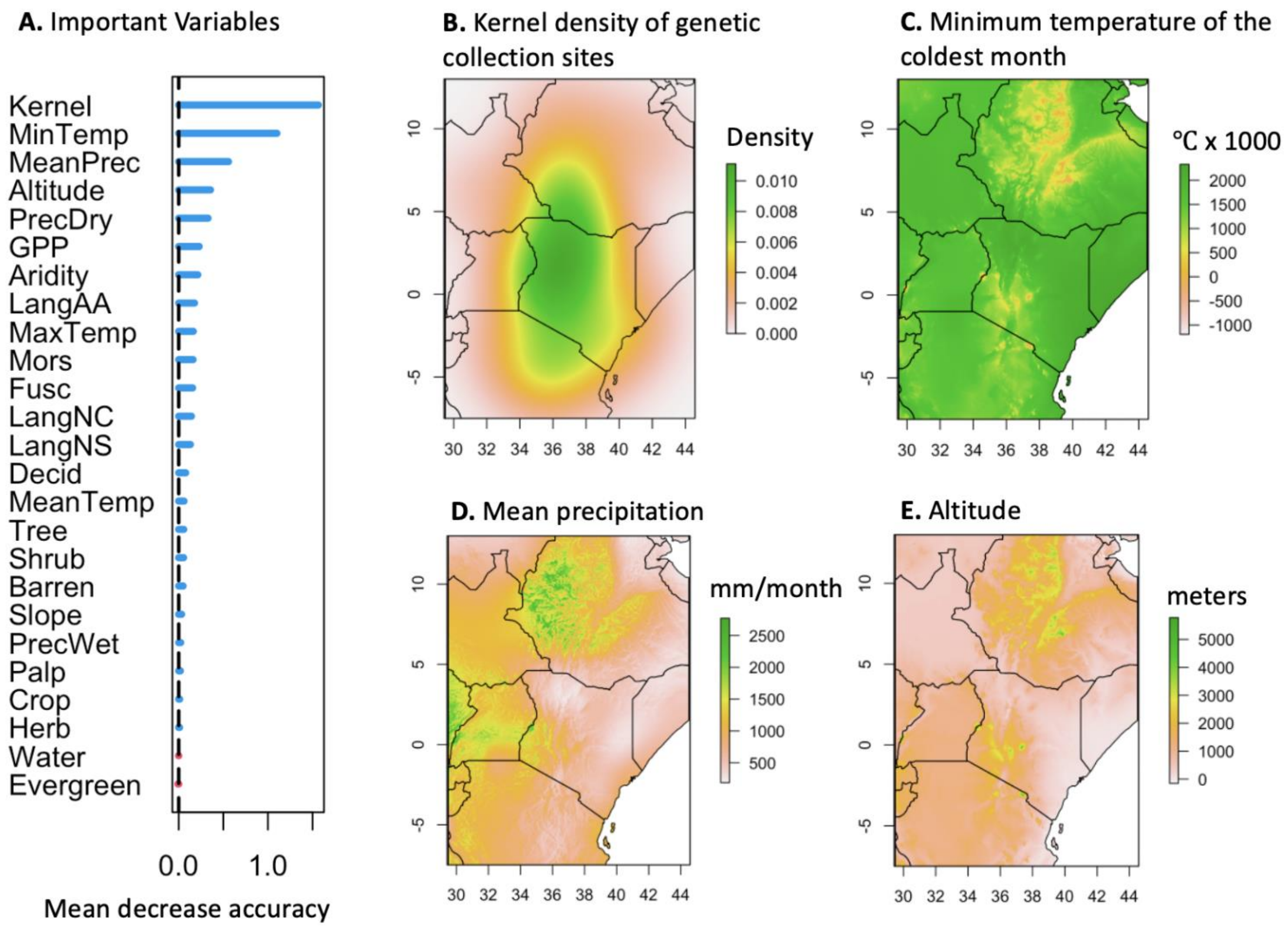


**Figure S2**: A. Most important variables in the SPRUCE random forest model for 2-4cM IBD tracts (~56 generations ago), determined by mean decrease in accuracy of the model when excluding each variable. B-E. Rasters of the four most important variables from this model run, overlaid with the country lines from eastern Africa.

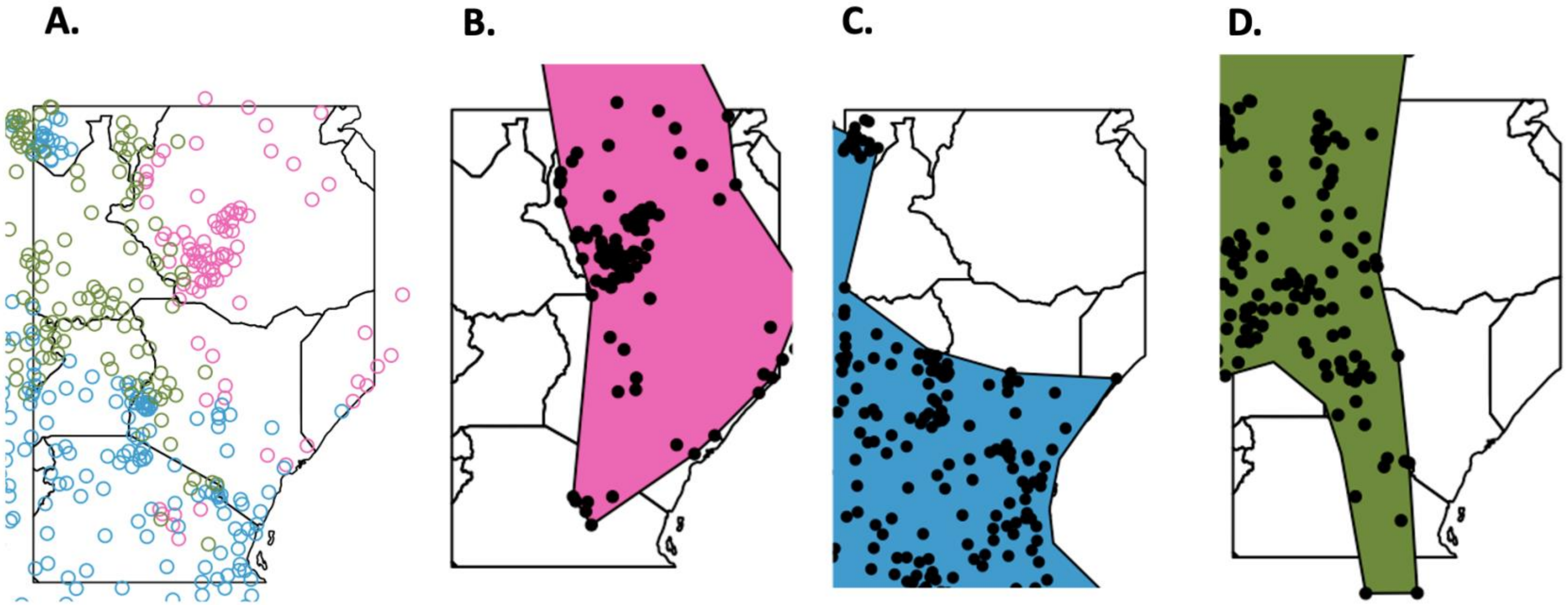


**Figure S3**: Creation of language family spatial variables. A. Location of the language family coordinates downloaded from Ethiolang and Wikitongues (pink = Afro-Asiatic, blue = Niger-Congo, green = Nilo-Saharan). B-D. Local convex hulls were created for each language family and then merged into one polygon to create categorical presence/absence variables. The final polygons are shown here for each language family, with the coordinates for each language family shown in black.

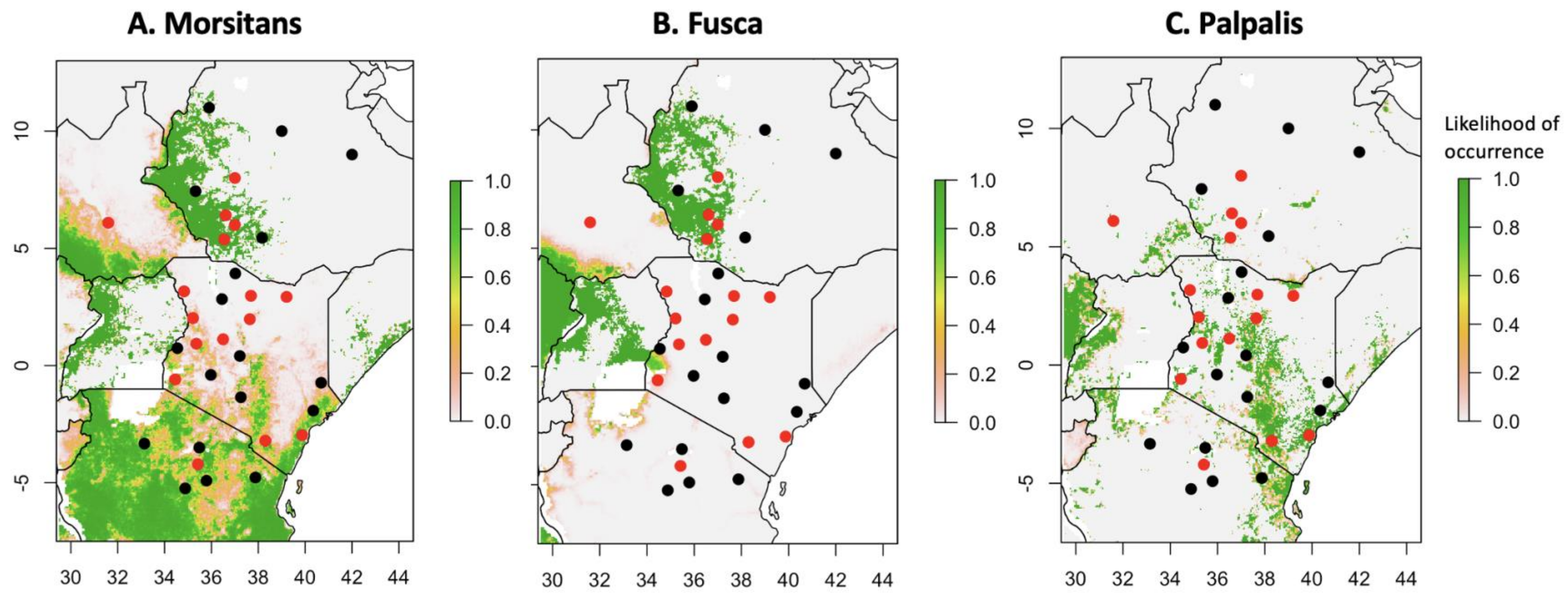


**Figure S4:** Tsetse suitability maps for the three main group (Morsitans, Fusca, and Palpalis) provided by the Food and Agriculture Organization of the United Nations (Cecchi 2002) and converted to rasters using qGIS 2.18 (QGIS Development Team 2017). Briefly, the morsitans group transmits *T. rhodensiense* in east Africa, the fusca group includes several vectors of livestock trypanosomiasis, and the palpalis group transmits *T. gambiense* in west and central Africa (Bouteille and Dumas 2003). Genetic sampling locations are also shown; points in red practice or recently practiced pastoralism or agropastoralism.

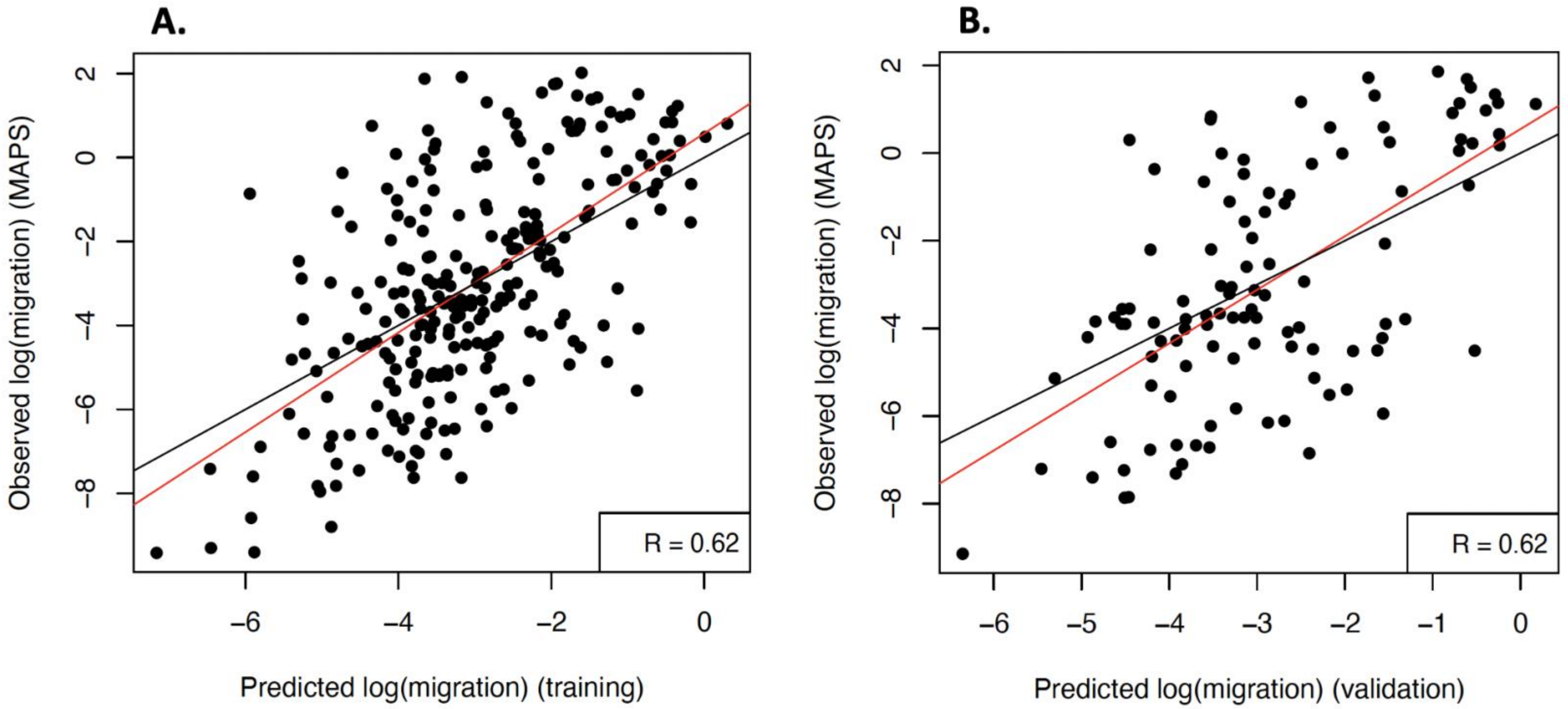


**Figure S5**: Observed migration (inferred by MAPS) versus predicted migration rate by the SPRUCE model for the training dataset (A.) and validation dataset (B.), suggesting the model is not overfitting the data. The random forest regression was trained on the full dataset for the oldest time period (~56 generations ago). The red line is the best-fit linear regression, and the black line is y=x.

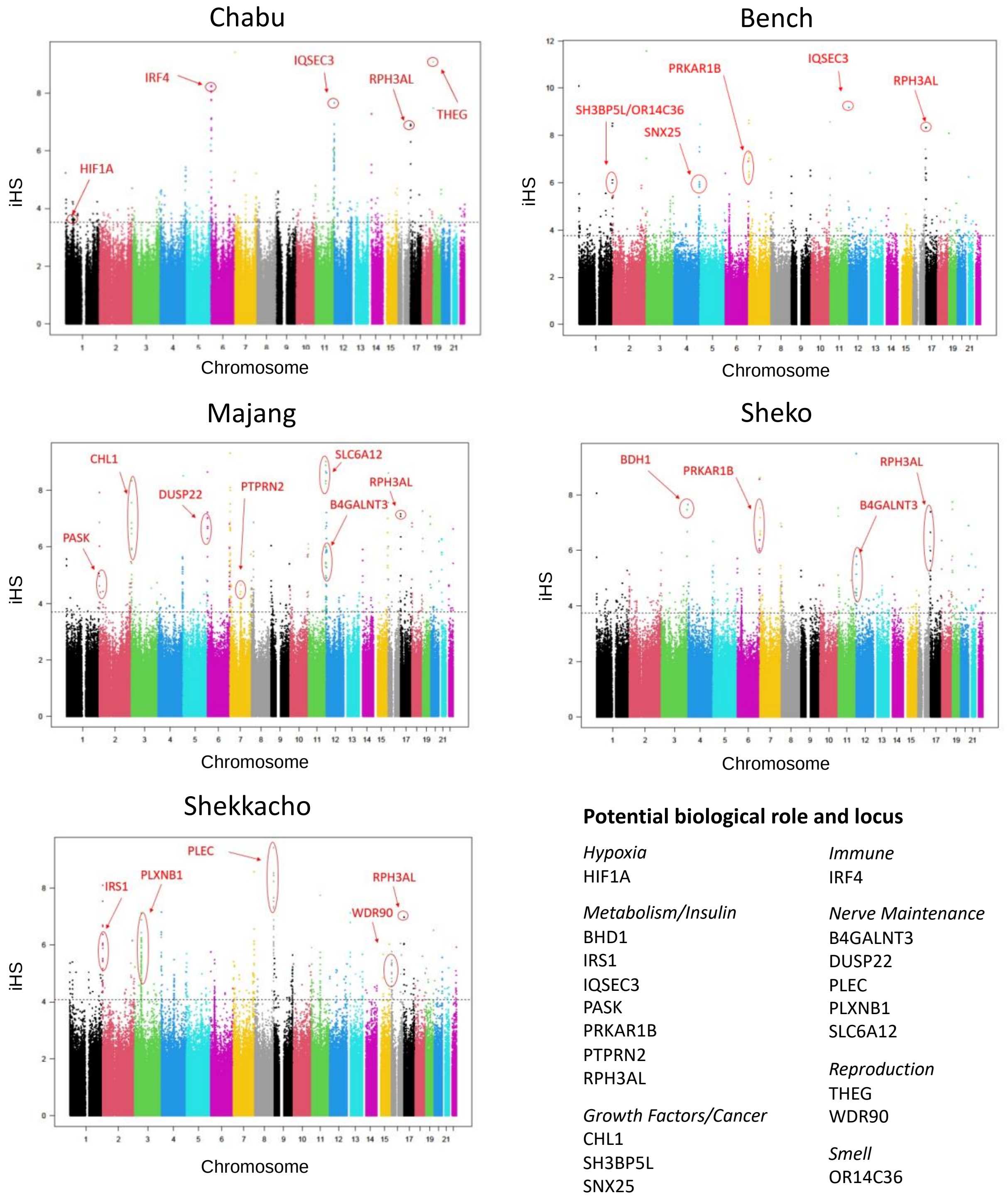


**Figure S6**: Selection scan results showing absolute value of integrated Haplotype Score (iHS) (Voight et al. 2006) across the genomes for five populations from southwestern Ethiopia. Higher values indicate higher likelihood of positive selection, based on decay of alleles from a query locus. The selection scan was performed with selscan 1.3.0 (Szpiech and Hernandez 2014).

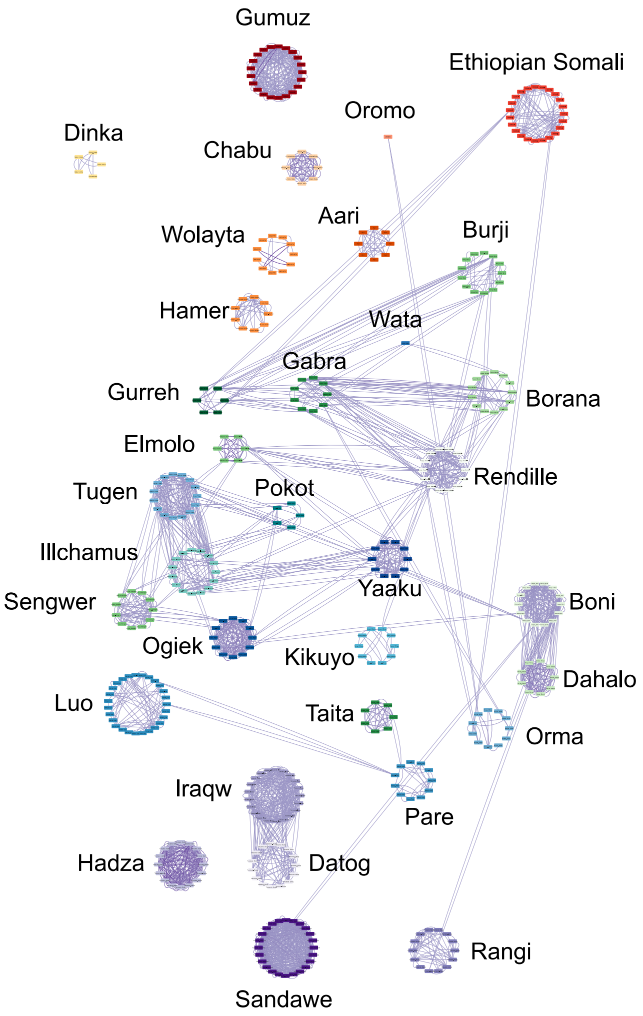


**Figure S7:** Identical by descent network created in Cytoscape (Shannon et al. 2003). Each individual is a node, and individuals from the same population are arranged in a circle. The locations of the circles correspond to the populations' approximate geographic locations. Edges between the nodes represent the number of shared IBD segments >6cM; however, edges representing <2 segments are not shown. Darker edges shows higher number of shared IBD segments. Individuals who do not share IBD segments with any other individuals are not shown.

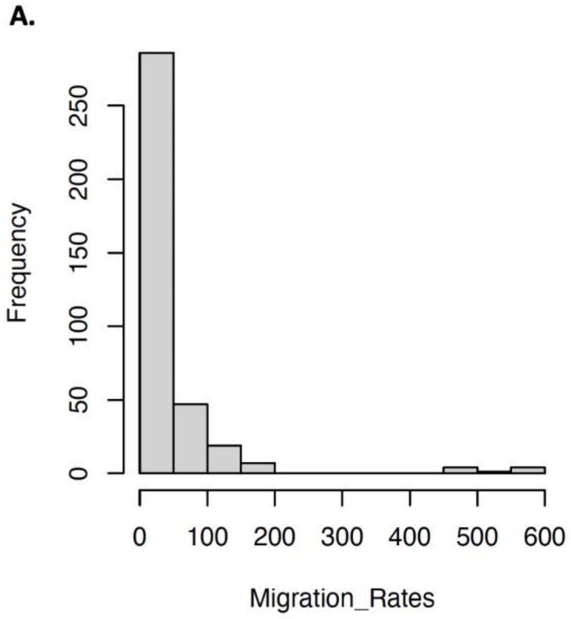


**Figure S8**: A. Histogram of migration rates from run 1 of MAPS using 2-4cM shared IBD tracts. Migration rates are unbalanced and skewed toward low values. B. By log transforming the migration rates, we obtained a more normal distribution to use as the observed variable in the SPRUCE model.

**Dataset S1 (separate file).** Candidate genes under positive selection in each of the five southwestern Ethiopia populations. chromosome, position, and for. For each candidate gene, chromosome, position, rsID, and iHS score are also listed (Voight et al. 2006). Candidate genes were determined by taking the top 0.1% largest absolute value iHS scores for each population.
